# Experiencing the wild: Red fox encounters are related to stronger nature connectedness, not anxiety, in people

**DOI:** 10.1101/2025.03.25.645298

**Authors:** F. Blake Morton, Carl D. Soulsbury

## Abstract

Global declines in biodiversity and human health are linked to declining human-nature connectedness, i.e., people’s subjective sense of their relationship with nature. Frequent, positive wildlife experiences can strengthen nature connectedness, improving public health and pro-environmental behaviours to protect biodiversity. Red foxes are the most widespread terrestrial carnivore on the planet. Urbanisation is associated with bolder fox behaviour and reduced public tolerance of foxes due to human-fox conflict. It remains unclear how interactions with foxes might influence (positively or negatively) nature connectedness and health in people, such as general anxiety. We surveyed 230 people using an online questionnaire. Nature connectedness was lower in urban areas and positively related to the perceived frequency and quality of fox encounters. Frequent, positive experiences were related to better attitudes and tolerance towards foxes, but unrelated to general anxiety. Management of human-fox coexistence should consider the potential role of foxes in shaping people’s broader nature connectedness.

## Introduction

Our planet is currently facing a biodiversity crisis (Ceballos & Ehrlich, 2017; Ceballos et al., 2015; Western, 1992), with future loss of species estimated up to 10,000 times higher than the expected rate (Pimm et al., 1995). At the same time, humanity is experiencing a global health crisis (Halder et al., 2023; Ney, 2012). Conditions such as obesity, diabetes, and heart disease have surged alongside doubling rates of adolescent depression and anxiety since the COVID-19 pandemic (Hossain et al., 2024; Powell-Wiley et al., 2021; Racine et al., 2021). Healthcare systems, such as the National Health Service (NHS), are under intense pressure due to long waiting lists, threatening their sustainability and infrastructure (Khan, 2023). These dual crises – biological and societal – highlight the urgent need for innovative and integrated solutions to improve global ecological stability and human well-being.

The accelerating pace of global urbanisation (Angel et al., 2011; Grimm et al., 2008; Nations, 2019) and the widespread decline in human-nature connectedness (hereafter ‘nature connectedness’) (Soga & Gaston, 2016, 2023), defined as a person’s subjective sense of their relationship with nature (Barragan-Jason et al., 2022), are key contributors to the biodiversity and human health crises. Studies reveal that nature connectedness is deteriorating globally at an alarming rate, driven by people spending less time in nature (Soga & Gaston, 2016). By 2050, most of the world’s human population will reside in urban settlements (Nations, 2019). Urbanisation diminishes natural habitats, limiting opportunities for people to engage physically, emotionally, and cognitively with nature (Derdouri et al., 2025; Hu et al., 2024). This disconnection from nature has profound consequences on people’s mental and physical health. Randomized controlled trials, for instance, show that experiences in nature improve nature connectedness (Barragan-Jason et al., 2022) and lead to improved cognition and lower levels of depression, rumination, and anxiety (Bratman, Daily, et al., 2015; Bratman, Hamilton, et al., 2015; Browning et al., 2023; Owens & Brunce, 2022). Randomized controlled trials also show that improving nature connectedness through experiences in nature can encourage people to behave pro-environmentally, such as living more sustainably or supporting conservation initiatives (Hsia et al., 2025; Mackay & Schmitt, 2019). Prioritising efforts to enhance nature connectedness in people, including urban residents, may therefore represent a targeted approach to address the biodiversity and health crises simultaneously (Soulsbury & White, 2019).

One promising avenue to foster deeper nature connectedness is the frequency and quality of human-wildlife interactions (Duke & Soulsbury, 2021). People who have more frequent and positive wildlife experiences, such as listening to birdsong and observing wild animal behaviour, show improved general health and stronger connections to nature (Chavez et al., 2023; Cox & Gaston, 2016; Cox & Gaston, 2018; Li et al., 2025). However, conflicts arising from close human-wildlife interactions have the potential to undermine the impact of these positive experiences (Chavez et al., 2023; Cox & Gaston, 2018). Greater personal anxiety among residents, for example, has been linked to increases in human-wildlife conflict in cities (Basak et al., 2022). Thus, frequent and positive interactions with wildlife must be balanced with managing negative experiences arising from human-wildlife cohabitation, including conflicts such as property damage, noise disturbances, and personal safety concerns (Nyhus, 2016; Soulsbury & White, 2015).

Interactions with species labelled as “pests” or “nuisances” are an example of the delicate balance between human-wildlife coexistence and conflict; negative experiences with these animals can shape negative attitudes and public intolerance (Hobson et al., 2024; Soulsbury & White, 2019), diminishing important factors related to public health, such as feelings of helplessness, anxiety, or unease (Basak et al., 2022; Zahl-Thanem et al., 2020). Managing human-wildlife conflict through effective education and other mitigation strategies may reduce the impact of negative experiences, potentially amplifying the positive effects of human-wildlife interactions (Konig et al., 2020). However, interactions with wildlife can be deeply subjective (Gillingham & Lee, 2003). Psychological biases, such as confirmation bias, generalisation, anthropomorphism, and framing effects, all contribute to how people choose to perceive interactions with other species (Ashish et al., 2022; Ballejo et al., 2021; Marsh & Hanlon, 2007; Waytz et al., 2014). Individuals may consider, for example, a fleeting glimpse of a rare species as profoundly meaningful, while others may only consider frequent encounters as significant. Similarly, one person’s positive experience with wildlife might be another person’s negative or neutral experience due to differences in emotion and expectations about the situation, including happiness, fear, or perceived risk (Castillo-Huitron et al., 2020; Johansson et al., 2025; Zahl-Thanem et al., 2020). Thus, subjective experiences, particularly in terms of their perceived frequency (e.g., rare versus often) and quality (e.g., positive versus negative), are important for understanding the potential role of human-wildlife interactions in shaping nature connectedness and public health among people.

Carnivores, particularly mesocarnivores, have learned to coexist remarkably well alongside humans due to their behavioural and ecological flexibility (Bateman & Fleming, 2012). Often appealing to ecotourists (van der Meer et al., 2016), wild carnivores are becoming increasingly prominent in urban landscapes (Bateman & Fleming, 2012), potentially offering opportunities to strengthen nature connectedness through public appreciation and engagement with nature (Giergiczny et al., 2022). Their charisma, coupled with their ecological importance (Ripple et al., 2014; Roemer et al., 2009), also makes carnivores potentially valuable for helping to maintain biodiversity more generally, improving opportunities for people to experience nature. However, interactions with wild carnivores also present important challenges due to the impact of people’s subjective experiences with them. Indeed, carnivores’ adaptability can lead to public nuisances, ranging from low-level disturbances (e.g., scent marking) to major health and safety risks (e.g., physical attacks) (Bombieri et al., 2023; Brand & Baldwin, 2020; Soulsbury & White, 2015). Such conflicts are often heavily sensationalised and framed negatively within popular culture (Arbieu et al., 2021; Bombieri et al., 2018; Bridge & Harris, 2020; Cassidy & Mills, 2012; Hughes et al., 2020; Nanni et al., 2020). This can distort public understanding and attitudes about the frequency and nature of those interactions (Casola et al., 2020). These subjective experiences can, in turn, influence public attitudes towards carnivores, leading to public intolerance (Brand & Baldwin, 2020; Kimmig et al., 2020). However, whether this subjectivity necessarily plays a role in shaping nature connectedness and public health among people remains uncertain due a lack of studies examining these relationships within the same population of people.

Red foxes (*Vulpes vulpes*) represent an important case study for understanding the relationships between nature connectedness, human health, and subjective wildlife experiences. As the most widely distributed wild carnivore on the planet (Marsh et al., 2022; Soulsbury et al., 2010), foxes have adapted to cohabitation with humans in a wide variety of urban and rural settings (Contesse et al., 2004; Plumer et al., 2014; Scott et al., 2014; Soulsbury et al., 2010). In some countries such as the UK, they are one of the most “liked” and frequently sighted mammals in people’s gardens (Baker et al., 2020). Their global distribution, positive public perception, and conspicuousness within neighbourhoods therefore suggests that people’s interactions with foxes are a potential gateway to the natural world, influencing nature connectedness and public health more generally (Soulsbury & White, 2019).

Foxes, however, also have a long and complex history of human persecution, leading to their public portrayal as a “nuisance” species (Padovani et al., 2021). For centuries, people have hunted foxes for sport and to protect livestock and game birds from being attacked or eaten (Jiguet, 2020; Kammerle et al., 2019; Porteus et al., 2019). In many cultures, foxes are heavily anthropomorphised, often described in stories as being “sly” or “cunning” in their interactions with people and other animals (Bridge & Harris, 2020; Harris & Baker, 2001; Morton et al., 2023). More recently, urbanisation has changed fox behaviour, with urban populations displaying greater boldness than rural populations (Gil-Fernandez et al., 2020; Morton et al., 2023). Increased boldness likely contributes to human-fox conflict, such as noise disturbance and damage to gardens (Brand & Baldwin, 2020), potentially explaining why positive attitudes towards foxes are reduced by as much as 50% in major cities such as London (Brand & Baldwin, 2020). Bolder fox behaviour is also heavily sensationalised within the media. Stories about physical attacks on pets and children, for instance, are taken out of context (Bridge & Harris, 2020; Cassidy & Mills, 2012; Harris & Baker, 2001; Scott et al., 2023), resulting in calls to cull urban foxes within these communities (Dowling, 2013). Such communications potentially fuel public safety concerns about foxes more generally (Brand & Baldwin, 2020; Bridge & Harris, 2020; Cassidy & Mills, 2012). To date, however, the relationships between public attitudes, tolerance, and the frequency and quality of human-fox interactions remains poorly understood due to a paucity of studies (Baker et al., 2004). It is also unknown whether, or how, subjective fox experiences might influence broader nature connectedness and health in people, particularly their general anxiety.

To address these gaps, we investigated the relationships between nature connectedness, anxiety, and people’s attitudes, tolerance, and subjective experiences with wild red foxes in the UK. Given the growing body of literature highlighting the positive impacts of human-wildlife interactions on nature connectedness and human health, we hypothesised that more frequent and better-quality fox experiences would predict greater nature connectedness and reduced anxiety in people. We also hypothesized that these subjective experiences would be related to people’s attitudes and tolerance towards foxes.

## Methods and materials

### Ethical statement

The study was approved by the Faculty of Health Sciences Ethics Committee of the University of Hull (FHS 24-25.042) and followed the research guidelines of the British Psychological Society.

### Participant recruitment

An online anonymous questionnaire of public anxiety, human-nature connectedness, and people’s experiences and attitudes towards foxes was administered to members of the UK general public in winter 2023 and 2025. The questionnaire was designed and managed using JISC software. Participants who were 18 years or older and a resident of the UK were recruited by posting an advert on social media and by circulating it to people and organisations (e.g., universities, museums, city councils, and wildlife groups) in London, Northeast England, Northwest England, Yorkshire and the Humber, East Midlands, East of England, Southeast England, and Scotland. To avoid disclosing the main purpose of the study to potential participants, the advertisement informed readers that the study was about “public health and environmental attitudes”.

### Questionnaire design

The questionnaire was composed of 29 questions (Supplementary Materials). Participants could not return to previously answered questions to change their responses. Although there was no time limit, the entire questionnaire should have taken no longer than 5-10 minutes to complete. The first question asked participants whether they gave consent prior to completing the questionnaire, and they were instructed that they could opt out of participating at any point before submitting their scores by simply closing the browser window.

#### Demographic questions

Three questions were demographic questions, asking for the participant’s age in years, educational background (e.g., GCSE’s or BSc), and whether they lived in an urban, rural, or suburban setting. “Urban” was defined as cities or densely populated areas. “Suburban” was defined as less densely populated areas built up around the outside of cities. “Rural” was defined as being further away from cities than suburban areas, with lots of undeveloped land. The questionnaire in 2025 also asked about the participant’s gender and in what region of the UK they lived.

#### General anxiety scale

Ten questions asked the participant to rate their level of anxiety within the last two weeks using a 7-point Likert scale, where 1=never and 7=all of the time. These questions were drawn from an existing, validated inventory for general anxiety (Levenstein et al., 1993). A composite score for each participant was derived from adding the ratings across questions (maximum of 70 points), where a higher score indicated a higher level of overall anxiety.

#### Human-nature connectedness scale

Ten questions asked the participant to rate, on a 7-point Likert scale, the extent to which they agreed to a series of statements related to personal attitudes and beliefs about nature, where 1=strongly disagree and 7=strongly agree. These questions were drawn from the *Connectedness to Nature Scale* – a validated and widely-used inventory for human-nature connectedness (Mayer & Frantz, 2004). A composite score for each participant was derived by adding the ratings of the eight questions that were “pro” environmental and subtracting this score by the sum of the scores from the two inverse questions. A maximum score of 53 was possible, with a higher score indicating a higher level of nature connectedness.

#### Public attitudes and tolerance towards foxes

Participants were shown an image of a red fox, followed by a 7-point Likert question asking how much the participant was willing to tolerate living alongside them, ranging from “very willing” to “very unwilling”. The participant was then asked to rate their overall attitude towards foxes, ranging from “very positive” to “very negative”. As discussed, public attitudes and tolerance are related but distinct constructs: Attitudes towards wildlife refer to people’s overall beliefs, feelings, and evaluations of wildlife, which are often characterised as being positive (e.g., valuing a species) or negative (e.g., seeing a species as dangerous) (Manfredo et al., 2009). Tolerance of wildlife, by contrast, refers to people’s willingness to coexist with another species, particularly in situations where tolerance may cause an inconvenience to a person, such as economic losses or physical risks (Zimmermann et al., 2020). Tolerance is influenced by attitudes, but also by other factors, such as societal norms and government legislation (Woolaston et al., 2021).

#### Subjective frequency and quality of fox experiences

The final section of the questionnaire asked the participant to rate, on a 5-point scale, their subjective quality of experiences with foxes, ranging from “very positive” to “very negative”. They were also asked to rate, on a 5-point scale, their subjective frequency of experiences with foxes, ranging from “very often” to “never”. Since subjective experiences with foxes were the primary focus of our study, we intentionally avoided giving the participant standardized definitions and metrics for “frequency” and “quality” to ensure that they applied their own subjective interpretations to these questions.

### Statistical analysis

We first tested how respondent age and education level influenced their nature connectedness and anxiety using linear models. Based on these results, we tested whether nature connectedness or general anxiety were related to age, urbanisation (rural, suburban, urban), and either the quality or frequency of experience with foxes as fixed effects. In each model, we included ‘urbanisation’ as an interaction and frequency/quality of fox experiences with education level included as a random effect. We finally tested whether attitude or tolerance towards foxes was related to age, urbanisation (rural, suburban, urban), and either the quality or frequency of experience with foxes as fixed effects. Again, we included urbanisation as an interaction and frequency/quality of fox experiences with education level included as a random effect. Significance of fixed effects in linear models were tested using a likelihood ratio χ^2^ test. The significance of the fixed effects was evaluated using type II Wald χ^2^ test in the R package *car* (Fox & Weisberg, 2018). LMMs were run using the *lmerTest* package (Kuznetsova et al., 2017), with all analysis carried out in R 4.3.1 (R Core Team 2023).

### Results Participants

A total of 230 people completed the questionnaire. Ages ranged between 18 to 83 years old (mean: 40 ± 16.7 years). Participants represented a variety of educational backgrounds, genders, habitat types, and region of the UK (Table 1). There was no overall effect of education (LR χ^2^= 0.10, p=0.748) or age (LR χ^2^=7.28, p=0.12) on nature connectedness. Instead, age (LR χ^2^=16.18, p<0.001; Figure S1), but not education (LR χ^2^=4.47, p=0.358), was related to general anxiety.

**Table 1.** Summary of participant demographics.

| Demographic Variable | % of participants |
| --- | --- |
| **Education** |  |
| No education | 1.3% |
| GCSE's | 7.8% |
| A levels / College diploma | 22.6% |
| BSc degree | 35.7% |
| Postgrad degree | 32.6% |
| **Gender** |  |
| Female | 62% |
| Male | 35% |
| Non-binary | 3% |
| **Habitat type** |  |
| Urban | 30.4% |
| Suburban | 41.3% |
| Rural | 28.2% |
| **Place of Residence** |  |
| Yorkshire and the Humber | 47% |
| Scotland | 21% |
| London and the southeast | 12% |

### The effect of urbanisation and subjective fox experiences on nature connectedness and anxiety

Nature connectedness was negatively related to urbanisation (Table 2, Figure 1) and positively related to frequency and quality of fox experiences (Table 2, Figure 2). Nature connectedness was unrelated to age or the interaction between urbanisation and fox experiences (Table 2). General anxiety was related to age (Table 2), but unrelated to urbanisation, the frequency and quality of fox experiences, or the interaction between urbanisation and fox experiences (Table 2).

**Figure 1.**
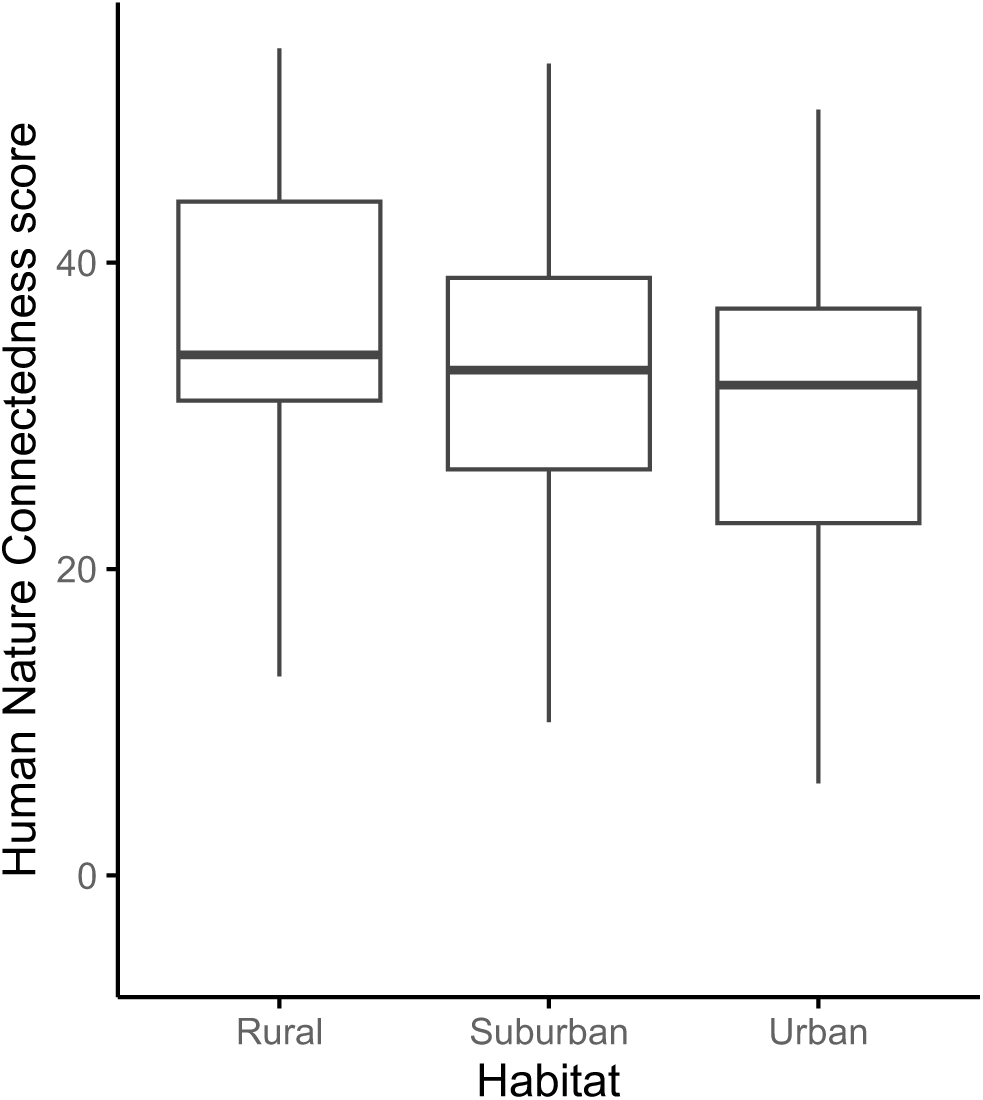
Median (+/- IQR) human-nature connectedness score in relation to urbanisation.

**Figure 2.**
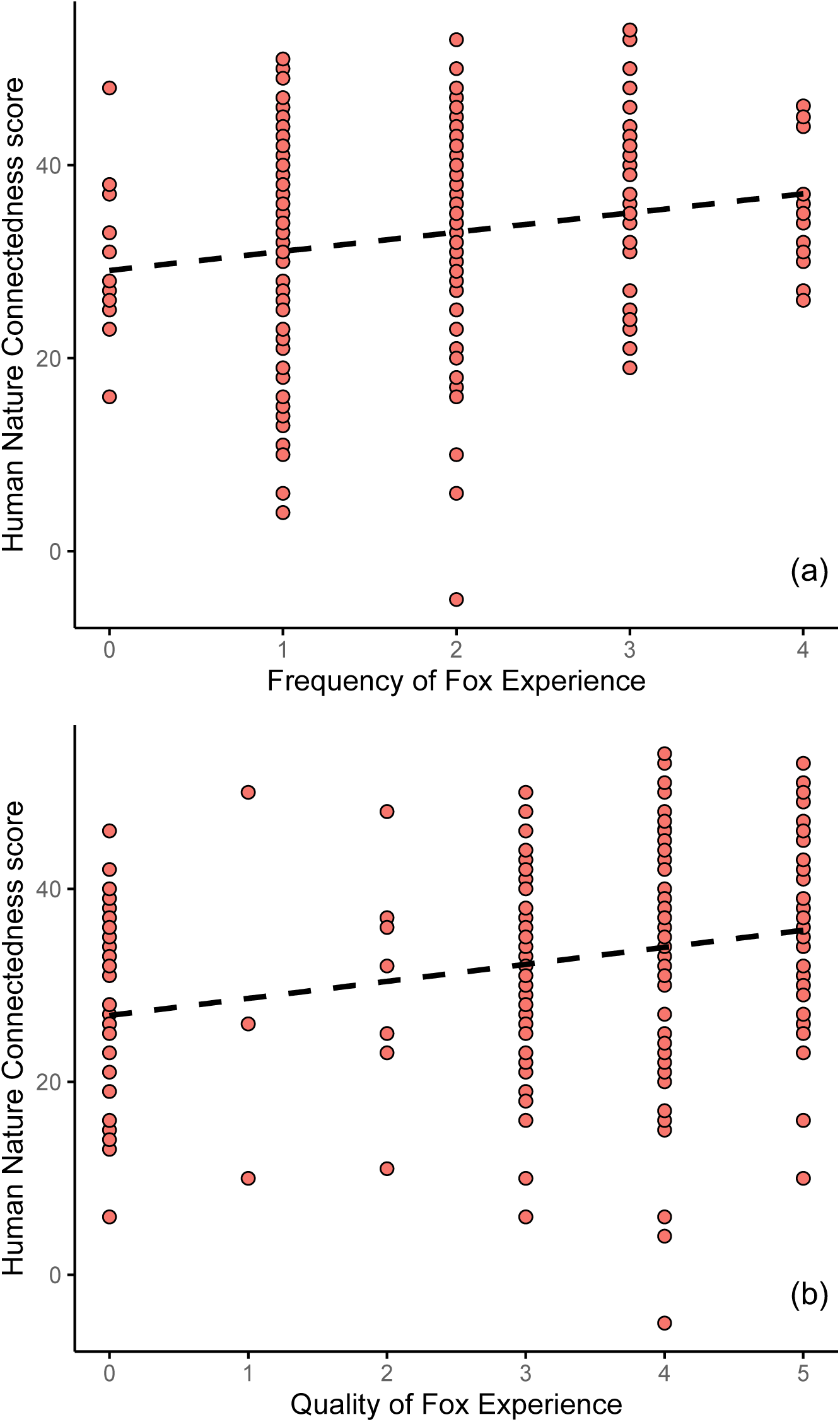
Relationship between participants’ human-nature connectedness scores and their (a) perceived frequency and (b) perceived quality of fox experiences.

**Table 2.** Nature connectedness (HNC) and general anxiety in relation to age, urbanisation, and the quality or frequency of fox experiences.

| Model parameter | HNC |  | General anxiety |  |
| --- | --- | --- | --- | --- |
| | Wald $\chi^2$ | *p* | Wald $\chi^2$ | *p* |
| Age | 0.31 | 0.580 | **11.48** | **<0.001** |
| Urbanisation | 4.74 | 0.094 | 2.18 | 0.336 |
| Quality | **14.42** | **<0.001** | 0.27 | 0.599 |
| Urbanisation x Quality | 0.49 | 0.783 | 2.45 | 0.294 |
| Age | 0.62 | 0.428 | **11.91** | **<0.001** |
| Urbanisation | **6.16** | **0.045** | 2.13 | 0.344 |
| Frequency | **7.74** | **0.005** | 1.91 | 0.167 |
| Urbanisation x Frequency | 2.53 | 0.283 | 0.18 | 0.914 |

### The effect of subjective fox experiences on public attitudes and tolerance

There was no effect of age or urbanisation on public attitudes or tolerance towards foxes (Table 3), but the quality of fox experiences was positively associated with both tolerance and attitudes (Table 3, Figure 3a and 3b). Likewise, the frequency of fox experiences was positively associated with tolerance and attitudes (Table 3, Figure 3c and 3d). There was no association between general anxiety and attitude or tolerance towards foxes (Table 3).

**Figure 3.**
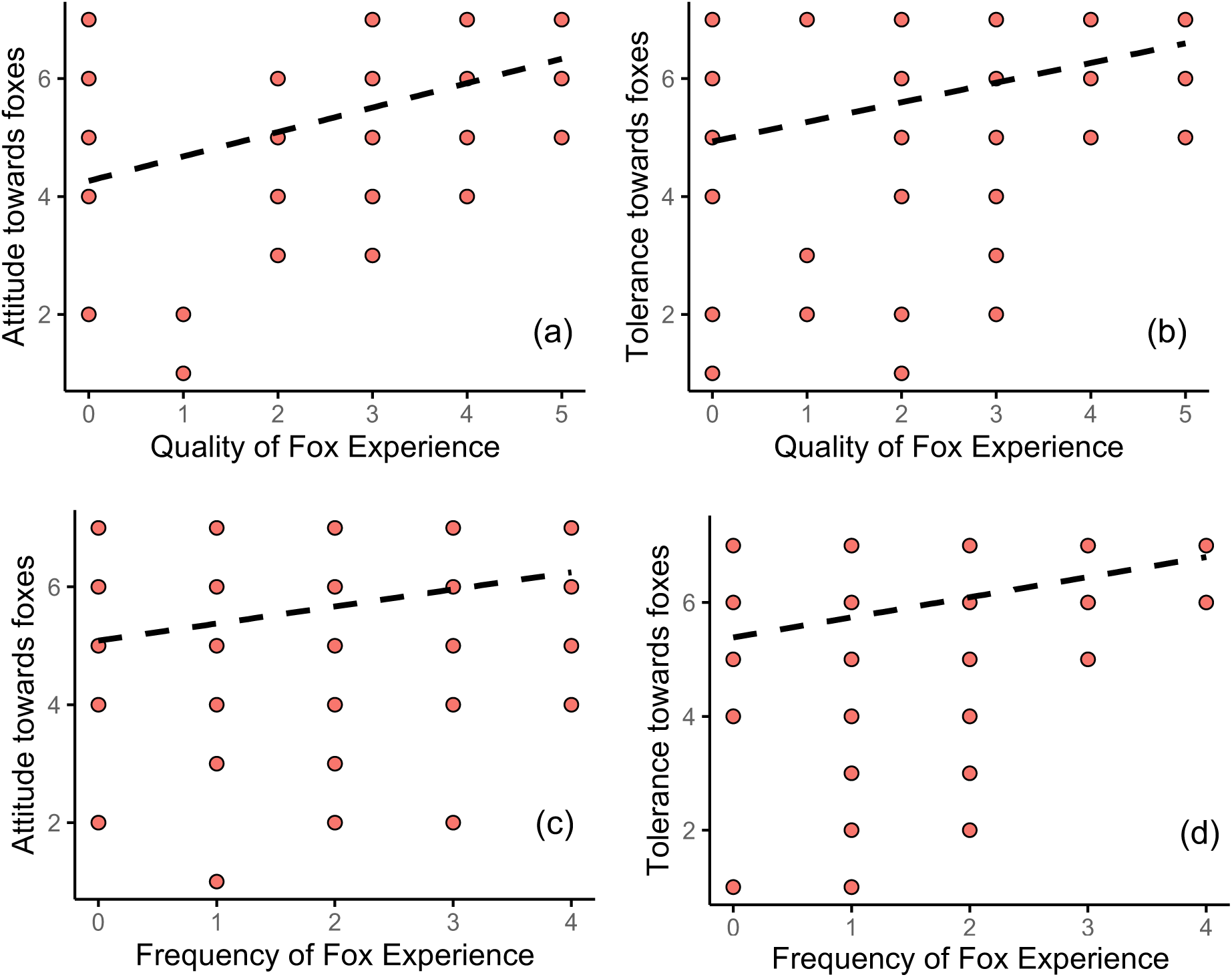
Relationship between perceived quality of fox experiences and participants’ (a) attitudes and (b) tolerance towards foxes, and the relationship between perceived frequency of fox experiences and participants’ (c) attitudes and (d) tolerance towards foxes.

**Table 3.** Attitudes and tolerance towards foxes in relation to age, urbanisation, and the frequency or quality of fox experiences.

| Model parameter | Attitude to foxes |  | Tolerance of foxes |  |
| --- | --- | --- | --- | --- |
| | Wald $\chi^2$ | *p* | Wald $\chi^2$ | *p* |
| Age | 0.11 | 0.736 | 0.02 | 0.882 |
| Urbanisation | 1.64 | 0.442 | 0.92 | 0.630 |
| Quality | **73.56** | **<0.001** | **45.17** | **<0.001** |
| Urbanisation x Quality | 1.45 | 0.483 | 0.842 | 0.656 |
| Age | 0.06 | 0.802 | 0.21 | 0.649 |
| Urbanisation | 1.95 | 0.377 | 1.81 | 0.405 |
| Frequency | **10.49** | **0.001** | **17.17** | **<0.001** |
| Urbanisation x Frequency | 0.42 | 0.809 | 0.35 | 0.841 |
| Age | 0.001 | 0.994 | 0.002 | 0.989 |
| Urbanisation | 1.86 | 0.394 | 1.73 | 0.422 |
| General anxiety | 0.41 | 0.523 | 1.53 | 0.217 |
| Urbanisation x Anxiety | 0.84 | 0.656 | 0.32 | 0.854 |

## Discussion

We investigated the relationships between nature connectedness, general anxiety, and people’s attitudes, tolerance, and subjective experiences with wild red foxes. We hypothesised that more frequent and better-quality experiences with foxes would be associated with greater nature connectedness and reduced anxiety in people. We also hypothesised that subjective fox experiences would be related, at least in part, to people’s attitudes and tolerance towards foxes. We found partial support for these hypotheses: perceived frequency and quality of fox experiences were positively related to public attitudes and tolerance towards foxes, as well as broader human-nature connectedness, but were unrelated to people’s level of anxiety.

Public attitudes, tolerance, and perceived frequency and quality of fox experiences were all related in the current study. Most people reported having positive attitudes and experiences with foxes, consistent with recent national surveys from the UK (Baker et al., 2020; Morton et al., 2024). People with more positive attitudes towards foxes were more tolerant of them in our study, and people with more positive attitudes and greater tolerance were also more likely to report having more frequent and positive fox experiences than other participants. These latter findings are consistent with Baker et al. (2004), which found that people were more likely to tolerate and develop positive relationships with foxes (e.g., feeding them) because they frequently saw them in their gardens. Collectively, in light of our findings and the global distribution, familiarity, and conspicuousness of foxes within neighbourhoods, further research should investigate whether positive interactions with foxes represent an important gateway to encourage broader nature connectedness (Methorst et al., 2020).

Frequency and quality of experiences with foxes were related to public attitudes and tolerance, suggesting that how people interpret these experiences (i.e., their attitudes towards foxes) is potentially just as important as the experiences themselves. Positive attitudes towards foxes may, therefore, potentially be important for promoting human-fox coexistence and strengthening people’s connection to nature. People with stronger connections to nature, for instance, might tolerate and feel more positively towards foxes, and therefore seek greater opportunities to interact with them (Cox & Gaston, 2018). They may also interpret their interactions with foxes more positively. Alternatively, frequent and more positive experiences with foxes may lead to greater tolerance and more positive attitudes, strengthening nature connectedness (Cox & Gaston, 2016; Cox & Gaston, 2018). Further work is needed to test these and other possible scenarios, ideally using longitudinal and randomly controlled trials to determine the direction and causality of the relationships identified in this study.

The negative correlation between nature connectedness and urbanisation in the current study is consistent with other studies (Richardson et al., 2022; Soga & Gaston, 2016), showing that urban residents have diminished levels of nature connectedness compared to rural and suburban settings. This may stem from the fact that urbanisation, as previously discussed, reduces people’s opportunities to physically, cognitively, and emotionally connect with nature (Derdouri et al., 2025; Hu et al., 2024). However, we found no significant interaction between urbanisation, people’s perceived frequency and quality of fox experiences, and human-nature connectedness, suggesting that although urban residents may feel less connected to nature, this is unlikely due to differences in how they perceive their experiences with foxes compared to more rural or suburban residents.

Studies show that exposure to nature, including positive interactions with wildlife, can improve health outcomes related to anxiety (Bratman, Hamilton, et al., 2015; Chavez et al., 2023; Morita et al., 2007; Owens & Brunce, 2022). In Poland, respondents have indicated that human-wildlife conflict can lead to personal anxiety; this mainly comes from species such as wild boar, but foxes were also reported to cause anxiety for some respondents (Basak et al., 2022). In our study, we found no direct relationship between nature connectedness and general anxiety, or between general anxiety and subjective fox experiences. Age and general anxiety were negatively related, consistent with other studies (Chaudhary et al., 2024), but many of our participants’ experiences with foxes were relatively infrequent. This may have prevented us from detecting the existence of small effects from these experiences on people’s levels of anxiety. Alternatively, experiences with foxes, while significant for nature connectedness, may not significantly impact the more complex interaction with people’s health, which is shaped by many other aspects of people’s lives. Browning et al. (2021) found a tendency for authors to publish positive results from studies of the impact of nature on public health, suggesting that such relationships are more nuanced. Indeed, health improvements can arise even when people’s exposure to nature is relatively short (e.g., 10 min) (Bettmann et al., 2024), and human-animal interactions can reduce anxiety even in sample sizes comparable or similar to our study (Beetz et al., 2012). Thus, the impact of fox experiences on human health may depend on other factors. Future studies should explore these and other possibilities, ideally using randomly controlled trials and other measures of health beyond the measure of anxiety used in the current study.

As discussed, foxes are currently one of the most liked mammals in countries such as the UK (Baker et al., 2020; Morton et al., 2024), but urbanisation has led to a greater likelihood of foxes behaving more boldly within cities compared to the countryside (Morton et al., 2023). This behavioural shift has likely contributed to the strikingly polarised pattern of public attitudes towards foxes in large metropolitan areas such as London (Brand & Baldwin, 2020). More broadly, recent rankings place the UK as the lowest in Europe, and one of the lowest globally, in terms of nature connectedness (Richardson et al., 2022; Swami et al., 2024). Studies also illustrate that the UK is one of the most ecologically depleted countries in the world (Burns et al., 2023; Lin et al., 2018). Thus, there is an urgent need to explore strategies to improve nature connectedness, particularly in the UK. Within this context, we suggest that supporting responsible and positive human-fox interactions could represent one such strategy, but the potential role of foxes in shaping British residents’ broader nature connectedness warrants further research.

## Conclusions

Global declines in biodiversity and human health are linked to declining opportunities for people to connect with nature. Our study highlights the intricate relationships between nature connectedness, public health, and subjective experiences with wildlife, using wild red foxes as a case study. Although we found no significant relationship between nature connectedness and human anxiety in the current study, tolerance and positive attitudes towards foxes, as well as frequent and high-quality experiences with them, emerged as significant predictors of broader nature connectedness. Together, these findings suggest that managing human-fox coexistence should consider the potential role of foxes in shaping people’s broader nature connectedness, particularly in cities where opportunities to experience nature are reduced.

## Data Availability

All data are provided in the supplementary materials (Dataset S1).

## Acknowledgments

We thank the participants and people who helped circulate the advertisement for recruitment. F.B.M. thanks the University of Hull for supporting the *British Carnivore Project*, including the current study, and the BCP team for fruitful discussions on human-fox interactions.

