## Supplementary materials for "Experiencing the wild: Red fox encounters are related to stronger nature connectedness, not anxiety, in people"

**Supplementary results**

At total of 233 participants consented to take part in the questionnaire (133 people in 2023, 100 people in 2025). Two participants were excluded because they left over half of the anxiety questions blank. One participant was excluded because they left over half of the nature connectedness questions blank. One person did not fill in one of the anxiety questions, and so we replaced it with the average score of the remaining nine items. Two people did not fill in one of the questions about positive nature connectedness, and so we replaced it with the average score of the remaining seven pro-environmental items.

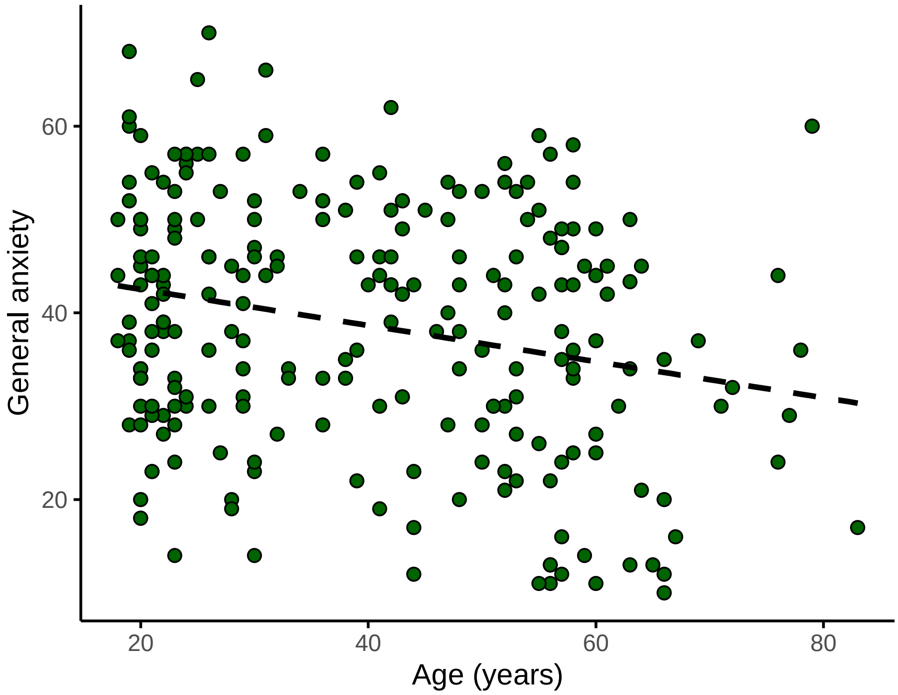


**Figure S1.** General anxiety in relation to participants' age.

**Consent form**

**Consent Form**

Please note that participation in this study requires that you consent to all following statements. If you decide to take part, please click "I consent to taking part". If you decide not to take part, please click "I do not consent" and submit the survey without responding to any questions.

- I confirm I have read the participation information sheet that explains the presented study.
- I have had the opportunity to consider the information and have been given contact details of the researcher to ask any questions or discuss the study.
- Any questions or concerns I have about the study have been answered satisfactorily.
- I understand that my participation is voluntary and that I am free to withdraw until the full submission of my answers without having to give a reason.
- I understand once I have completed the study, I cannot withdraw my anonymized data.
- I understand that the research data, which are not linked to me, will be retained by the researchers and may be shared with others and publicly disseminated to support other research in the future.
- I agree to take part in the presented study.

**1. Do you give your consent to take part in this study? ***

◯ I give consent to take part in the study

◯ I do not give consent to take part in this study

**Online questionnaire administered to human participants.**

Question 1. What is your age in years?
Question 2. What is your highest level of education? Please tick one.

- None
- GCSE's
- A-levels/college
- University degree (First year)
- University degree (Second year)
- University degree (Third year)
- Postgraduate qualification

Question 3. What is your gender?

Question 4. What region of the UK do you live?

Question 5. In what type of area do you live?

- Urban (cities or densely populated areas)
- Suburban (less densely populated areas built up around the outside of cities)
- Rural (further away from cities than suburban areas, lots of undeveloped land)

Question 6: On a scale of 1-7, how often have you experienced **feeling restless, keyed up, or on edge** in the past 2 weeks?

- 1 (Never)
- 2 (Rarely)
- 3 (Occasionally)
- 4 (Sometimes)
- 5 (Often)
- 6 (Very often)
- 7 (All the time)

Question 7: On a scale of 1-7, how often have you experienced **trouble relaxing** in the past 2 weeks?

- 1 (Never)
- 2 (Rarely)
- 3 (Occasionally)
- 4 (Sometimes)
- 5 (Often)
- 6 (Very often)
- 7 (All the time)

Question 8: On a scale of 1-7, how often have you experienced **muscle tension or tightness** in the past 2 weeks?

- 1 (Never)
- 2 (Rarely)
- 3 (Occasionally)
- 4 (Sometimes)
- 5 (Often)
- 6 (Very often)
- 7 (All the time)

Question 9: On a scale of 1-7, how often have you experienced **worrying** in the past 2 weeks?

- 1 (Never)
- 2 (Rarely)
- 3 (Occasionally)
- 4 (Sometimes)
- 5 (Often)
- 6 (Very often)
- 7 (All the time)

Question 10: On a scale of 1-7, how often have you experienced **anxiety** in the past 2 weeks?

- 1 (Never)
- 2 (Rarely)
- 3 (Occasionally)
- 4 (Sometimes)
- 5 (Often)
- 6 (Very often)
- 7 (All the time)

Question 11: On a scale of 1-7, how often have you experienced **irritability or grouchiness** in the past 2 weeks?

- 1 (Never)
- 2 (Rarely)
- 3 (Occasionally)
- 4 (Sometimes)
- 5 (Often)
- 6 (Very often)
- 7 (All the time)

Question 12: On a scale of 1-7, how often have you experienced **too many things to do** in the past 2 weeks?

- 1 (Never)
- 2 (Rarely)
- 3 (Occasionally)
- 4 (Sometimes)
- 5 (Often)
- 6 (Very often)
- 7 (All the time)

Question 13: On a scale of 1-7, how often have you experienced **feeling mentally exhausted** in the past 2 weeks?

- 1 (Never)
- 2 (Rarely)
- 3 (Occasionally)
- 4 (Sometimes)
- 5 (Often)
- 6 (Very often)
- 7 (All the time)

Question 14: On a scale of 1-7, how often have you experienced **feeling under pressure from deadlines** in the past 2 weeks?

- 1 (Never)
- 2 (Rarely)
- 3 (Occasionally)
- 4 (Sometimes)
- 5 (Often)
- 6 (Very often)
- 7 (All the time)

Question 15: On a scale of 1-7, how often have you experienced **problems that seem to be piling up** in the past 2 weeks?

- 1 (Never)
- 2 (Rarely)
- 3 (Occasionally)
- 4 (Sometimes)
- 5 (Often)

- 6 (Very often)
- 7 (All the time)

Question 16: On a scale of 1-7, how strongly do you agree with this statement:
**I often feel a sense of oneness with the natural world around me.**
- 1 (Strongly disagree)
- 2
- 3
- 4 (Neither agree nor disagree)
- 5
- 6
- 7 (Strongly agree)

Question 17: On a scale of 1-7, how strongly do you agree with this statement:
**I think of the natural world as a community to which I belong.**
- 1 (Strongly disagree)
- 2
- 3
- 4 (Neither agree nor disagree)
- 5
- 6
- 7 (Strongly agree)

Question 18: On a scale of 1-7, how strongly do you agree with this statement:
**I recognise and appreciate the intelligence of other living organisms.**
- 1 (Strongly disagree)
- 2
- 3
- 4 (Neither agree nor disagree)
- 5
- 6
- 7 (Strongly agree)

Question 19: On a scale of 1-7, how strongly do you agree with this statement:
**I often feel disconnected from nature.**
- 1 (Strongly disagree)
- 2
- 3
- 4 (Neither agree nor disagree)
- 5
- 6
- 7 (Strongly agree)

Question 20: On a scale of 1-7, how strongly do you agree with this statement:
**When I think of my life, I imagine myself to be part of a larger cyclical process of living.**
- 1 (Strongly disagree)
- 2
- 3
- 4 (Neither agree nor disagree)
- 5
- 6
- 7 (Strongly agree)

Question 21: On a scale of 1-7, how strongly do you agree with this statement:
**I often feel a kinship with animals and plants.**
- 1 (Strongly disagree)
- 2
- 3
- 4 (Neither agree nor disagree)
- 5
- 6
- 7 (Strongly agree)

Question 22: On a scale of 1-7, how strongly do you agree with this statement:
**I have a deep understanding of how my actions affect the natural world.**
- 1 (Strongly disagree)
- 2
- 3
- 4 (Neither agree nor disagree)
- 5
- 6
- 7 (Strongly agree)

Question 23: On a scale of 1-7, how strongly do you agree with this statement:
**Like a tree can be part of a forest, I feel embedded within the broader natural world.**
- 1 (Strongly disagree)
- 2
- 3
- 4 (Neither agree nor disagree)
- 5
- 6
- 7 (Strongly agree)

Question 24: On a scale of 1-7, how strongly do you agree with this statement:
**When I think of my place on earth, I consider myself to be a top member of a hierarchy that exists in nature.**
- 1 (Strongly disagree)
- 2
- 3
- 4 (Neither agree nor disagree)
- 5
- 6
- 7 (Strongly agree)

Question 25: On a scale of 1-7, how strongly do you agree with this statement:
**My personal welfare is dependent on the welfare of the natural world.**
- 1 (Strongly disagree)
- 2
- 3
- 4 (Neither agree nor disagree)
- 5
- 6
- 7 (Strongly agree)

**Please look at this picture**

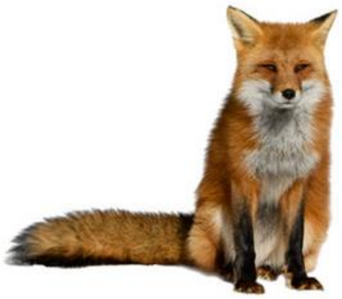

Question 26: How much are you willing to live alongside foxes? Please tick one.
- o Very willing
- o Willing
- o Somewhat willing
- o Neither willing nor unwilling
- o Somewhat unwilling
- o Unwilling
- o Very unwilling

Question 27: How do you feel towards foxes? Please tick one
- o Very Positive
- o Positive
- o Somewhat positive
- o Neutral (neither positive nor negative)
- o Somewhat negative
- o Negative
- o Very negative

Question 28: How would you describe your personal experience with foxes? Please tick one.
- o Very positive
- o Positive
- o Neutral
- o Negative
- o Very negative
- o I don’t have any personal experience with this animal

Question 29: How often do you encounter foxes? Please tick one.
- o Very often
- o Often
- o Sometimes
- o Rarely
- o Never
